# Accelerated sampling of protein dynamics using BioEmu augmented molecular simulation

**DOI:** 10.64898/2026.01.07.698041

**Authors:** Soumendranath Bhakat, Eva-Maria Strauch

## Abstract

We introduce a workflow that integrates BioEmu-generated conformational ensemble with physics-based molecular simulations and Markov State Models to sample Boltzmann-weighted conformational populations across biomolecules. Molecular simulations initiated from BioEmu ensemble capture active-to-inactive transitions in CDK2 and BRAF, two members of the serine-threonine kinase family, and elucidate how the disease-causing V600E mutation in BRAF drives population shifts among distinct metastable states relative to the wild type. Furthermore, we combined BioEmu ensemble with experimental cryo-EM data to construct all-atom conformational ensembles of biomolecules. In comparison to the AlphaFold2 reduced multiple sequence alignment (*rMSA-AF2*) approach, BioEmu-generated ensembles sample a broader conformational space for serine–threonine kinases but fail to capture conformational heterogeneity in several cases, including Glycine transporter 1 (GlyT1), a membrane transporter, and plasmepsin-II (PlmII), an aspartic protease. Systems where side-chain conformational heterogeneity governs protein dynamics such as cryptic pocket opening in PlmII or transitions between multiple metastable states in GlyT1; molecular simulations initiated from BioEmu generated ensemble do not capture the full spectrum of conformational heterogeneity. Overall, this study presents a straightforward framework for integrating generative AI based protein emulators with statistical physics to recover Boltzmann-weighted conformational ensembles at scale, while also highlighting critical limitations that necessitate careful, system-specific analysis when interpreting protein conformational landscapes.

## Introduction

Protein dynamics are central to biological function, and exploiting conformational motion to develop therapeutics is a cornerstone of dynamics driven drug discovery. Functional regulation in proteins is rarely governed by a single static structure; instead, it emerges from transitions between metastable states that modulates cryptic pocket opening, protein–protein interactions, and allosteric signal propagation. While high-resolution structures obtained from X-ray crystallography or cryo-electron microscopy provide invaluable snapshots of terminal states, such as the active and inactive forms of kinases and GPCRs, they do not capture the relative populations of these states or how environmental perturbations, such as ligand binding or disease-causing mutations shift these populations(1–5).

Molecular dynamics (MD) simulations provide a physics-based framework for probing the thermodynamics and kinetics of protein motion(6, 7). However, unbiased MD is fundamentally limited by the *timescale problem*: biologically relevant conformational transitions are often separated by large free-energy barriers that are inaccessible on practical simulation timescales(8). Enhanced sampling strategies have been developed to address this limitation and broadly fall into two classes: methods that bias along predefined collective variables(9–12), and methods that globally modify the potential energy surface through temperature scaling(13) or history-dependent bias potentials(14). While powerful, these approaches suffer from critical drawbacks, most notably their reliance on prior knowledge of reaction coordinates (collective variables) and the fact that the resulting population distributions are not intrinsically physical, requiring careful reweighting and specialized protocols to recover unbiased thermodynamics.

An alternative class of approaches, such as adaptive sampling, seeks to remain unbiased by iteratively launching ensembles of short MD trajectories from diverse conformational states, often identified via clustering of initial simulation data. While conceptually appealing, adaptive sampling workflows typically depend on carefully tuned clustering strategies, state definitions, and feedback protocols, and require specialized software frameworks and expert intervention(15, 16). As a result, these methods often exhibit a steep learning curve and limited accessibility, particularly when applied to complex and biologically relevant conformational transitions.

Recent advances in generative AI-based protein structure/ensemble prediction algorithms have raised the question of whether data-driven models can bridge this sampling gap(17–21). Methods based on AlphaFold2 (AF2), including AF2-RAVE(22, 23), AF2-MSM(24) and CryoPhold(25), exploit reduced multiple sequence alignments (rMSA) to induce conformational diversity. The central idea behind rMSA-AF2 is to generate a heterogeneous set of initial structures that can seed downstream unbiased MD simulations, thereby accelerating conformational exploration. While this strategy can capture cryptic pockets and alternate conformational states, the resulting physics refined ensembles remain strongly biased toward the “*coverage*” of the initial ensemble. If the initial coverage of rMSA-AF2 fails to capture meaningful diversity, it is unlikely that seeding short MD simulations will substantially enhance this sampling.

BioEmu(26) offers a fundamentally different paradigm. Rather than perturbing the inputs to a static structure predictor, BioEmu is a generative diffusion model fine-tuned on molecular dynamics simulation data and trained to reproduce statistically independent equilibrium structural distributions. This enables substantially *broader coverage* of conformational space compared to rMSA-AF2. However, BioEmu generated ensembles alone lack explicit physical populations and kinetic information. To recover physical meaning, we integrate BioEmu generated conformational ensemble with molecular simulation (MD) and Markov State Models (MSMs)(27–29), yielding a unified framework that provides quantitative state populations and statistically rigorous free energy landscapes of conformational transitions expressed along physically meaningful and human readable collective variables.

To date, the molecular simulation community has largely focused on variation of rMSA-AF2 seeded MD workflows, where conformational diversity is introduced by subsampling evolutionary information. In contrast, molecular simulations initiated from BioEmu generated conformational ensemble replaces MSA perturbations with generative sampling, enabling direct comparison of these two paradigms. In this work, we systematically evaluate BioEmu guided MD simulation against rMSA-AF2 augmented MD in their ability to capture functionally relevant conformational transitions. We focus on two serine/threonine kinases (CDK2 and BRAF), in which transitions between the *DFG-in* (active) and *DFG-out* (inactive) states involve large conformational barriers and are typically induced by ligand binding (30). This makes them an ideal system to assess whether such transitions can be sampled *a priori*, without ligands, biasing potentials, or prohibitively long simulations starting from a single experimental structure. We further demonstrate that molecular simulations initiated from BioEmu ensemble captures mutation-induced population shifts with significantly reduced cumulative simulation time compared to rMSA-AF2-based approaches(31).

Integrating single-molecule experimental techniques such as cryo-electron microscopy (cryo-EM) with generative AI based ensemble generator to construct experimentally informed conformational ensembles is an active area of research. In this work, we extend the CryoPhold framework(25), which integrates generative AI derived ensembles with experimental cryo-EM maps and physics-based molecular simulations; to incorporate BioEmu-generated conformational ensembles as an initial prior. We further analyze two distinct and challenging systems to examine the limitations of molecular simulations initiated from BioEmu with the rMSA-AF2 approach. Together, our results establish BioEmu-guided MD simulations as a straightforward protocol for accelerating access to protein motion beyond the reach of traditional enhanced sampling, enabling practical exploration of functionally important protein conformations central to drug discovery.

## Results

### BioEmu seeded molecular simulations captures conformational dynamics in kinases

The conformational transition between DFG-in and DFG-out states is a rare event in serine– threonine kinases such as CDK2 and BRAF and typically occurs only in the presence of type II inhibitors. This makes these kinases an ideal test case to assess whether BioEmu- or rMSA-AF2– seeded molecular simulations can capture these rare conformational transitions *a priori*.

Molecular simulations initiated from BioEmu generated conformational ensemble sampled the equilibrium free-energy landscapes of apo BRAF and CDK2, capturing conformational transitions between the DFG-in and DFG-out states. *rMSA-AF2* generated conformational ensembles predominantly restricted to the DFG-in state, whereas all-atom reconstruction of BioEmu generated ensembles populated a broader conformational space beyond the DFG-in state (**Figure 2 A-D**). Markov State Models constructed using two key structural variables capturing the DFG transition resolved not only terminal DFG-in and DFG-out states but also intermediate states (DFG_Neo_; Figure S1 in Supporting Information) and additional substates, key features of local conformational dynamics.

**Figure 1.**
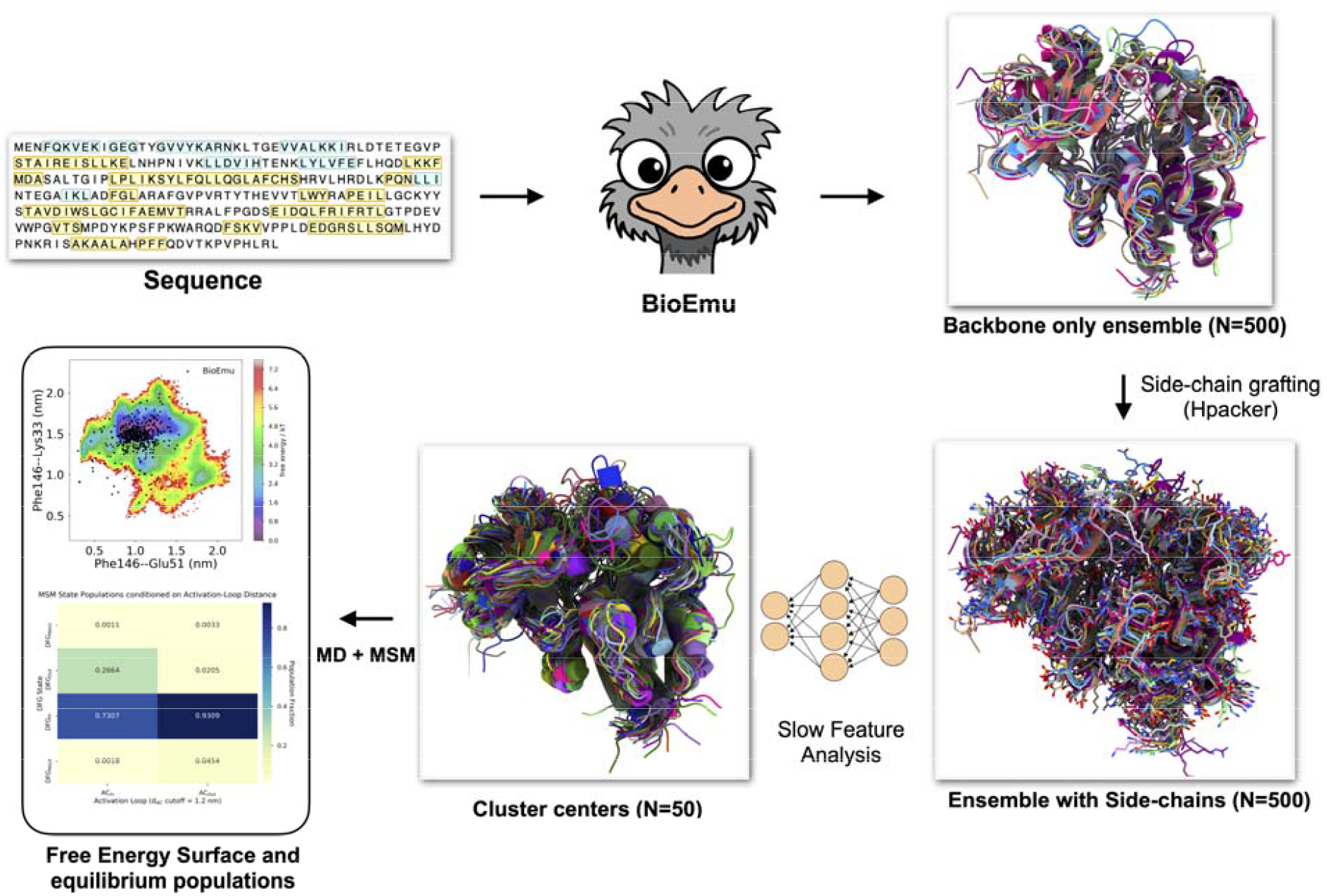
Schematic representation of BioEmu seeded molecular simulation workflow. BioEmu takes a protein sequence as input and generates a backbone only conformational ensemble of a protein monomer. Hpacker(32) is then used to add side chains and produce all-atom representations of the ensemble. Slow feature analysis is performed on Cα–Cα distances, and *K means* clustering (N = 50) on the first two slow features is used to generate a latent structural ensemble. These structures are used to initiate 50 independent 100 ns MD simulations for each protein system studied in this manuscript. The combined simulation data are used to build an MSM, from which MSM weighted free-energy surfaces are projected along functionally relevant, human-readable collective variables, and the populations of macrostates and substates are computed from the equilibrium distribution.

**Figure 2.**
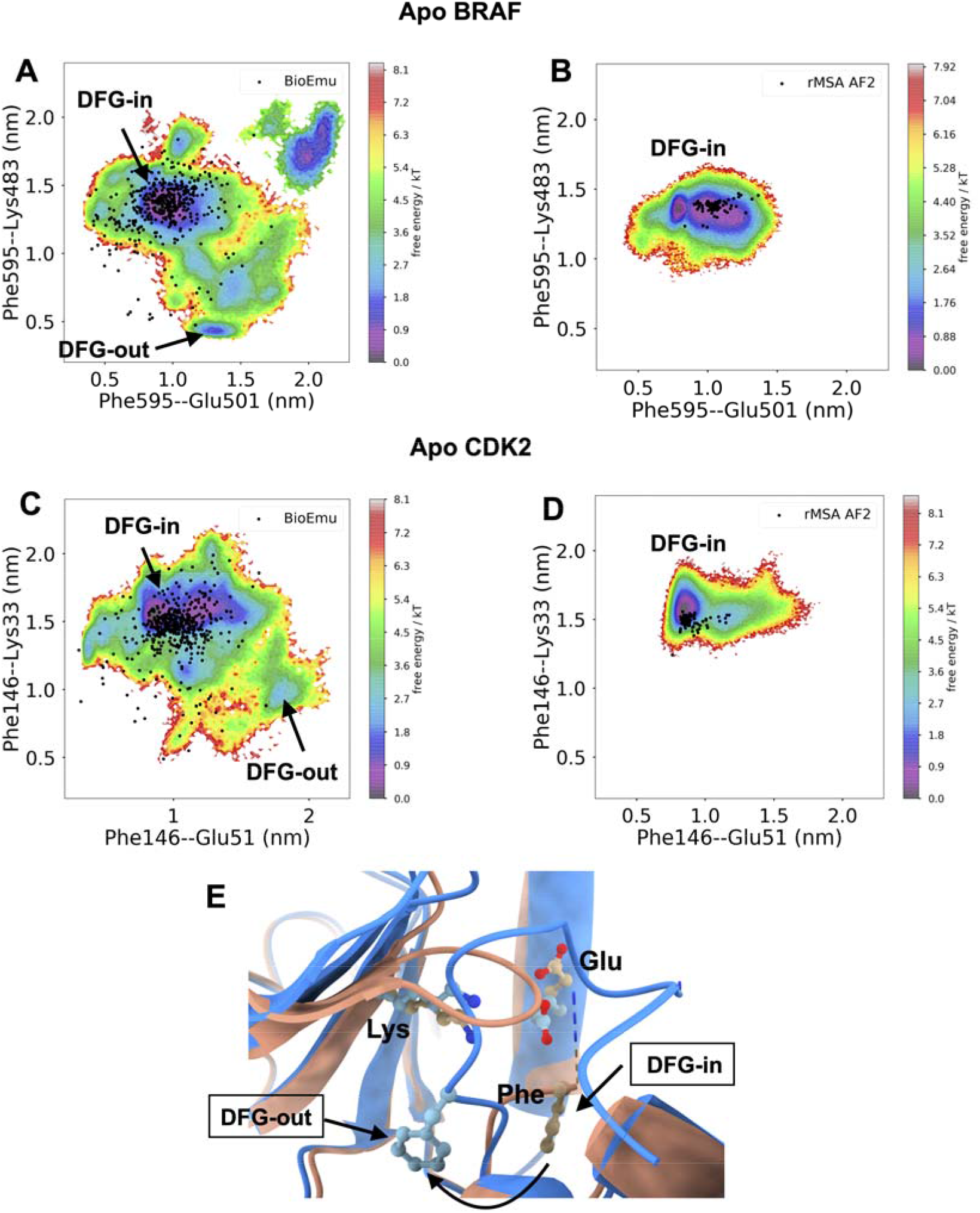
MSM weighted free-energy surfaces resolve DFG-in to DFG-out conformational transitions. MSM weighted free-energy surfaces projected along key inter-residue distances (Phe Cα–Glu Cα (x-axis) and Lys Cα–Phe Cζ (y-axis)) capture conformational transitions between DFG-in and DFG-out states. BioEmu seeded molecular simulations (total simulation time: 5 µs) sample the DFG-in to DFG-out transition in apo wild-type BRAF and CDK2 (A, C), whereas rMSA-AF2 augmented MD simulations (total simulation time: 8 µs) remain confined to the DFG-in state (B, D). Black dots denote conformational ensembles generated by BioEmu (n = 500) and rMSA-AF2 (n = 80), illustrating the broader conformational coverage achieved by BioEmu relative to rMSA-AF2. Representative structural orientations of the DFG motif are shown for the DFG-in (salmon) and DFG-out (cyan) states, highlighting the relative positioning of the DFG phenylalanine, lysine, and glutamate residues (E).

In apo CDK2, MD simulations started from BioEmu ensemble captures pronounced conformational heterogeneity within the DFG-in ensemble, with substates defined by the χ1 dihedral angle (Phe_N_, Phe_F1_, Phe_F2_) of the DFG-Phe. We further capture transitions between extended (AC_out_) and folded (AC_in_) activation-loop conformations and quantify their populations within both DFG-in and DFG-out macrostates (**Figure 3 A, B**). Transition from DFG-in to DFG-out shifts the activation-loop population from the extended AC_out_ state toward the folded AC_in_ state, a hallmark of the active-to-inactive conformational transition in apo CDK2 (33, 34). In parallel, the DFG-in to DFG-out transition is accompanied by a population shift of the *αC-helix* from the in (LG_L_) to the out (LG_U_) conformation. Together, the simultaneous capture of activation loop extended-to-folded and *α*C-helix in-to-out transitions highlights the ability of BioEmu augmented, physics-based molecular simulations to resolve the full spectrum of conformational heterogeneity underlying the active to inactive transition of apo CDK2.

**Figure 3.**
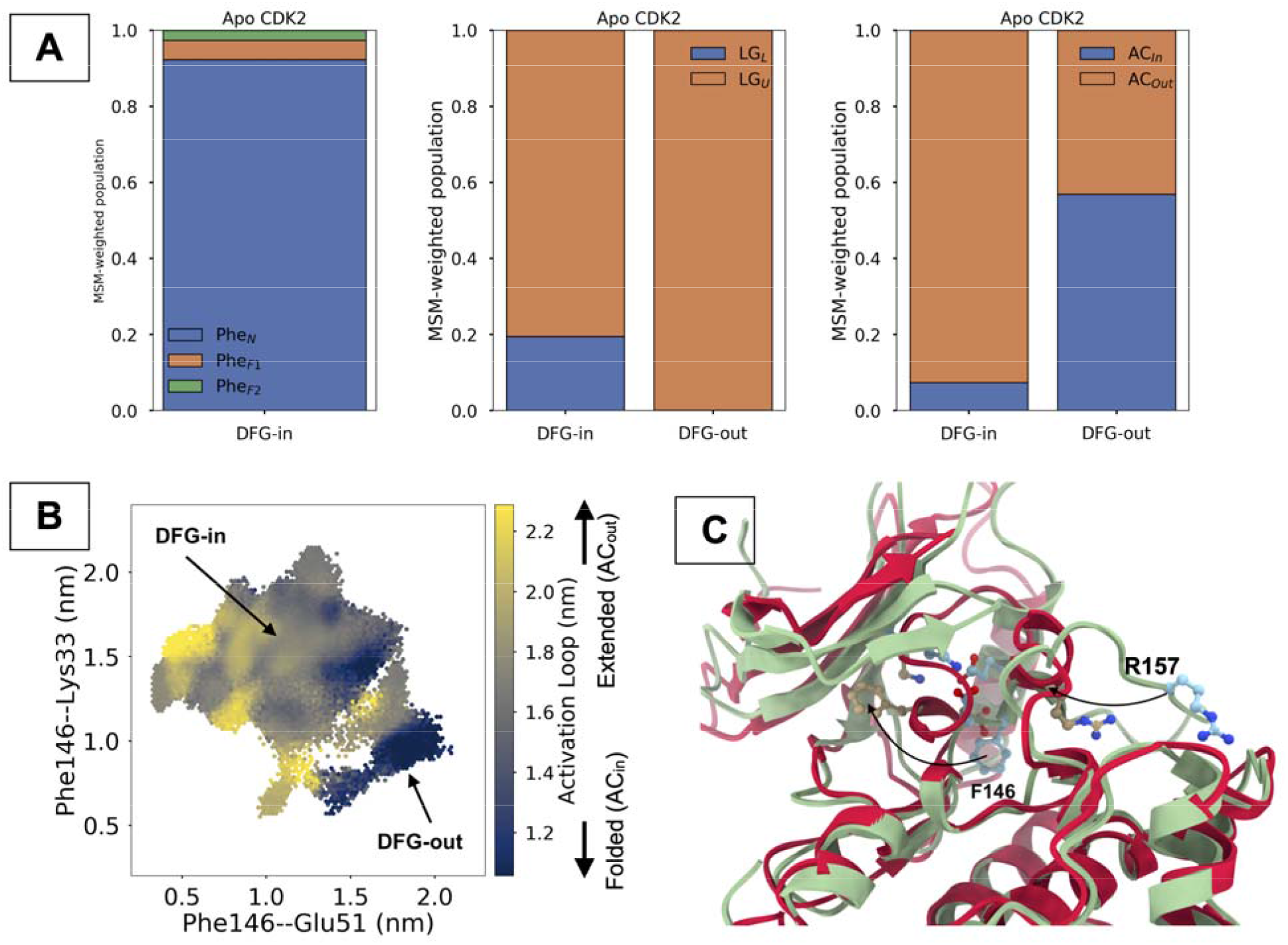
Normalized MSM-weighted populations of alternate DFG-Phe states (Phe_N_, χ1 ≈ −1 rad; Phe_F1_, χ1 ≈ + 1 rad; and Phe_F2_, χ1 ≈ ±π rad) within the DFG-in ensemble, together with the relative populations of the *α*C-helix in the “in” (LG_L_) and “out” (LG_U_) conformations and of the activation loop in extended (AC_out_) and folded (AC_in_) states, highlight the ability of BioEmu augmented molecular simulations to capture conformational heterogeneity in apo CDK2 (A). Projection of the activation-loop distance (D145-CA–R157-CA) onto two key distances distinguishing DFG-in and DFG-out states reveals a higher population of the folded activation loop in the DFG-out state compared to DFG-in (B). Superposition of representative DFG-in and DFG-out structures highlights structural changes, including flipping of the DFG-Phe side chain and folded activation loop, hallmark of active↔inactive transition (C).

We next investigated whether BioEmu augmented MD simulations capture mutation-induced population shifts between wild-type and V600E BRAF. The simulations successfully resolve a “*population shift*” of the DFG-Phe sidechain rotamer from the Phe_F1_ state (χ1 ≈ −1 rad) to the Phe_N_ state (χ1 ≈ +1 rad) within the DFG-in macrostate (**Figure 4**), a hallmark conformational feature associated with V600E-mediated BRAF activation (35, 36). In addition, the simulations capture population shifts of the *αC-helix* between the “*in*” (LG_L_, L=latched) and “*out*” (LG_U_, U=unlatched) conformations across DFG-in and DFG-out macrostates for both wild-type and V600E BRAF. Notably, the V600E mutation biases the ensemble toward the *α*C-helix “*in*” (LG_L_) conformation in both DFG-in and DFG-out states, consistent with the active apo BRAF conformational landscape induced by mutation.

**Figure 4.**
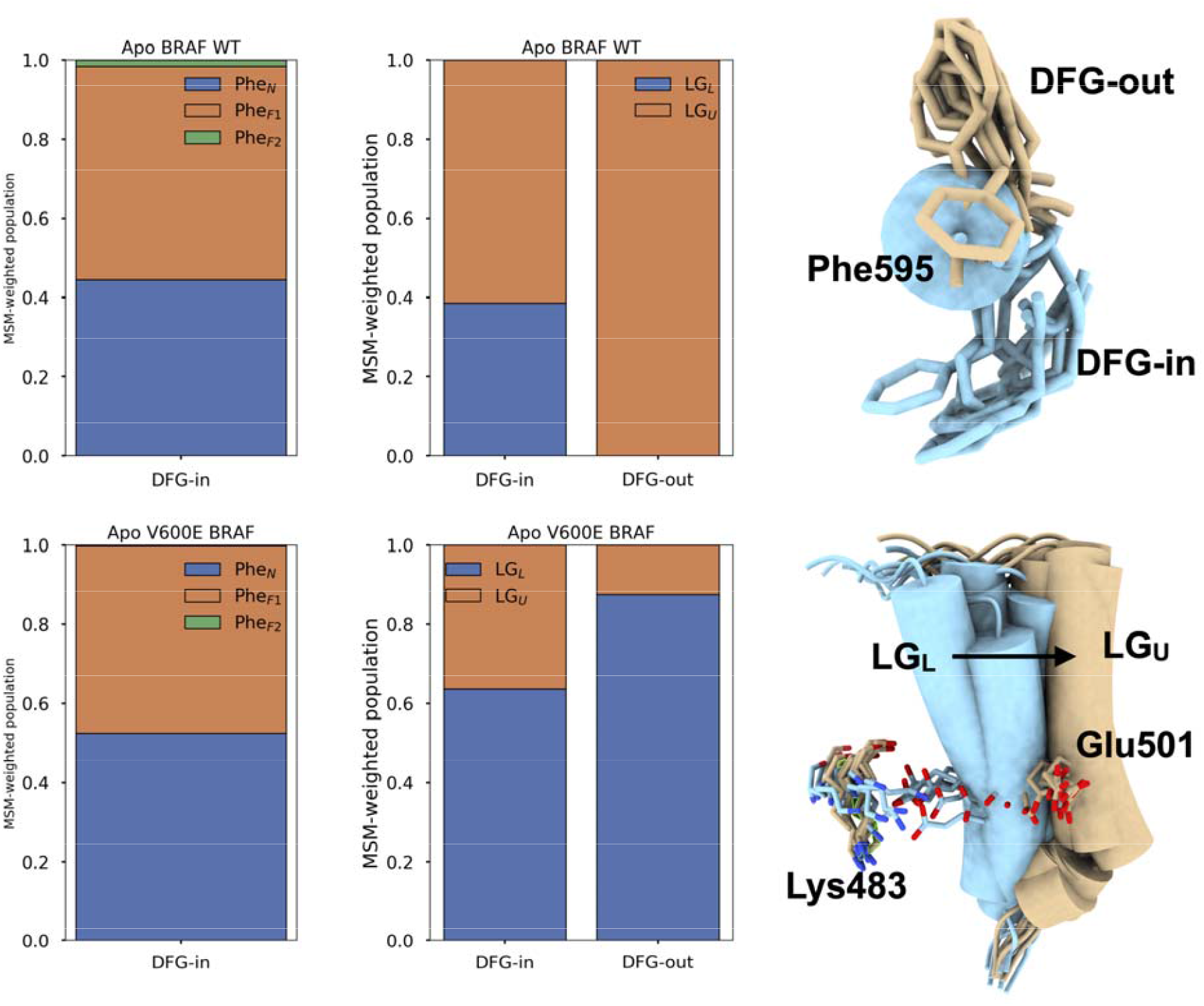
Normalized MSM-weighted populations of the Phe_N_ (χ1 ≈ −1 rad), Phe_F1_ (χ1 ≈ +1 rad), and Phe_F2_ (χ1 ≈ ±π rad) states within the DFG-in macrostate, together with the relative populations of the *α*C-helix in the “in” (LG_L_) and “out” (LG_U_) conformations across DFG-in and DFG-out states, highlight how the V600E mutation shifts the ensemble toward more active-like conformations.

### Integration of BioEmu with Cryo-EM and molecular simulation captures partial conformational transitions in GlyT1

Glycine transporter 1 (GlyT1) is a membrane transporter that modulates neurotransmission by clearing extracellular glycine and is a key drug target for the treatment of schizophrenia(37). GlyT1 adopts three distinct conformational states that are characteristic of many transporter proteins: (a) an *occluded* state bound to glycine (EMDB: EMD-37492; 2.58 Å), (b) an *inward* state bound to inhibitor, ALX-5407 (EMDB: EMD-37493; 3.35 Å), and (c) an *outward* state bound to inhibitors, SSR-504734 (EMDB: EMD-37494; 3.22 Å) and PF-03463275. Fitting a single structural model into a cryo-EM density does not provide information about the relative populations of these conformational states. To address this limitation, we recently introduced CryoPhold, a framework that combines generative AI based ensemble generation with experimental cryo-EM maps through Bayesian reweighting(25). Clustering of BioEmu-generated conformational ensembles followed by Bayesian reweighting against experimental cryo-EM maps (*CryoEmu*) enabled the generation of all-atom conformational ensembles of GlyT1 and estimation of their relative populations across inward, outward, and occluded states (**Figure 5**). Molecular simulations initiated from ensembles corresponding to each state and integrated using Markov state models (MSMs) enabled us to construct free-energy surfaces along key structural distances that describe transitions among inward, occluded, and outward states. Simulations initiated from CryoEmu generated ensembles successfully sampled the outward and occluded states but only partially sampled the inward state and the flipping of Y62, a key residue governing conformational transitions in GlyT1. In contrast, simulations initiated from rMSA-AF2 derived ensembles captured full *inward*↔*occluded*↔*outward* transitions as well as Y62 flipping(25). These results indicate that CryoEmu augmented simulations do not fully recover the conformational heterogeneity of GlyT1 and highlight that rMSA-AF2 generated ensembles provide broader coverage of the GlyT1 conformational landscape compared to BioEmu derived ensembles.

**Figure 5.**
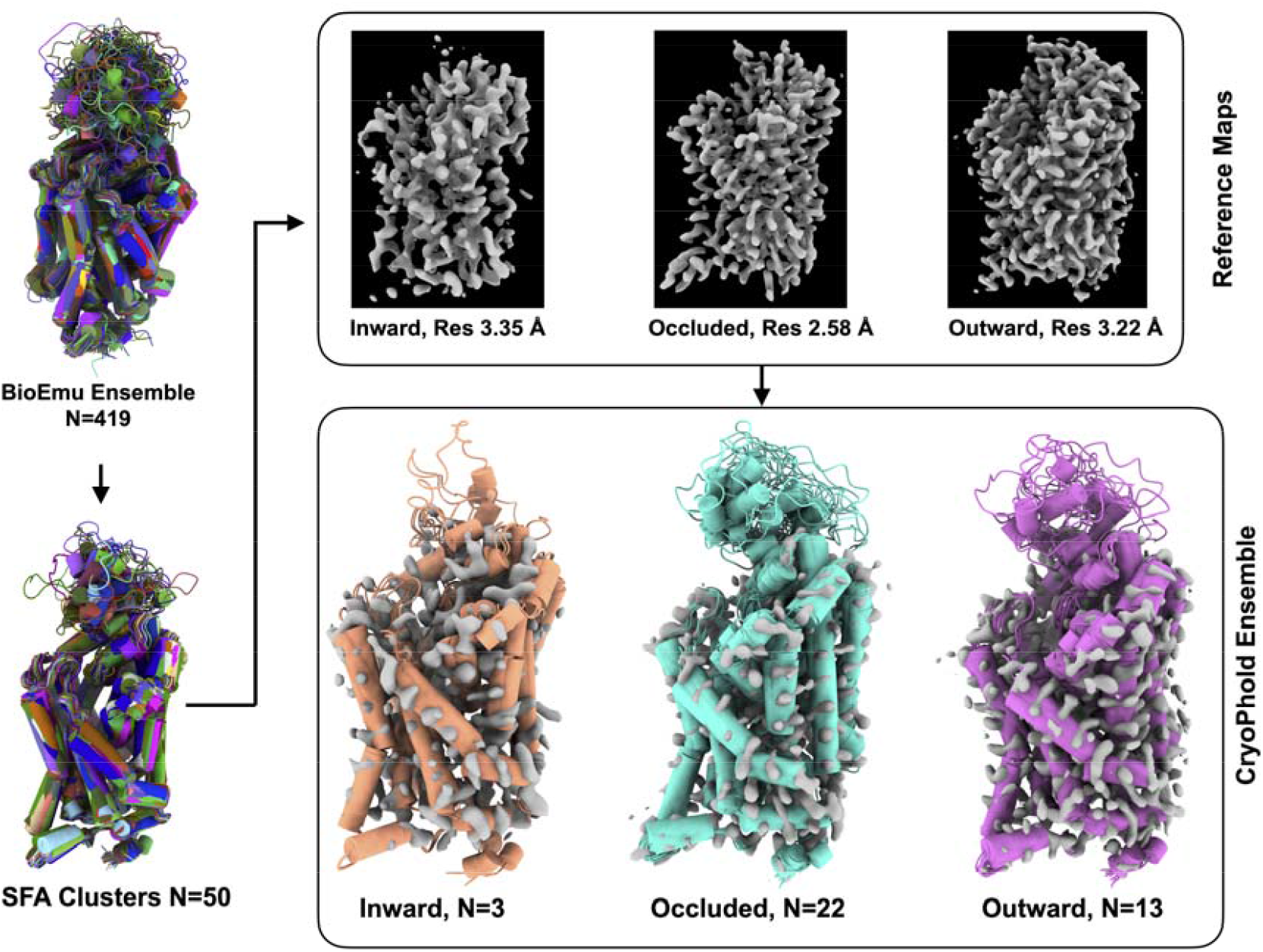
For apo GlyT1, we generated an initial conformational ensemble using BioEmu, which was subsequently subjected to SFA based clustering to obtain a reduced ensemble of 50 representative structures. Bayesian reweighting of this SFA derived ensemble using the CryoPhold workflow enabled the construction of all-atom conformational ensembles corresponding to the inward, outward, and occluded states of GlyT1. This combined workflow, referred to as CryoEmu, also provides an initial estimate of the relative populations of the alternative GlyT1 conformational states.

### BioEmu generated ensemble fails to capture sidechain conformations necessary for cryptic pocket opening in plasmepsin-II

Plasmepsin-II (PlmII) is an aspartic protease and a key drug target for malaria(38). *Meller* et al. previously used rMSA-AF2 augmented molecular simulations to capture cryptic pocket opening and closing in PlmII(24). A critical conformational motion regulating cryptic pocket opening in PlmII is the *flipping* of Trp41, making this system well-suited for evaluating the ability of the all-atom BioEmu ensemble to capture cryptic pocket opening. Comparison of BioEmu and rMSA-AF2 augmented molecular simulations demonstrates that the latter more effectively captures Trp41-mediated cryptic pocket opening in PlmII (**Figure 7**). This difference directly reflects the narrower coverage of Trp41 sidechain conformational heterogeneity in BioEmu generated ensemble compared to those produced using the rMSA-AF2 approach.

## Conclusions

Herein, we demonstrate how molecular simulations started from BioEmu generated conformational ensemble, resolve functionally important conformational states and mutation-induced population shifts beyond what is captured by rMSA-AF2 augmented molecular simulations. A key finding is that, for the serine-threonine kinases reported here, BioEmu-generated conformational ensembles provide substantially broader conformational coverage than rMSA-AF2-derived ensembles, a critical factor governing accelerated sampling when integrated with physics-based molecular simulation. However, conformations generated by BioEmu alone do not encode physical information, such as state populations or weights, making it challenging to prioritize functionally relevant states for downstream drug discovery workflows, such as population-weighted ensemble docking(39). By incorporating physics-based molecular simulations on top of the BioEmu-generated ensemble, we assign statistically meaningful weights to macro and microstates and capture mutation-induced population shifts, a hallmark of many human diseases. While initiating short MD simulations from approximately ∼500 BioEmu-generated structures could be computationally demanding, we mitigate this cost by training slow feature analysis (SFA)(39) on *C*α*–Cα* distances derived from the BioEmu ensemble and clustering along the first two slow features, yielding a reduced latent ensemble of 50 representative structures. For a protein of ∼200–500 residues, initiating ∼500 independent simulations of 100 ns each corresponds to an aggregate sampling of ∼50 μs. At typical performance of ∼200–300 ns/day per trajectory on an NVIDIA A100 GPU, each trajectory would require ∼0.3–0.5 days; thus, running all 500 simulations in parallel would require ∼500 A100 GPUs for ∼8–12 hours. In contrast, restricting simulations to 50 SFA-selected representative structures reduces the aggregate sampling to ∼5 μs and requires only ∼50 A100 GPUs for the same wall-clock time, reducing GPU demand by an order of magnitude while preserving coverage of the slow, functionally relevant conformational landscape. This reduction enables efficient parallel simulations on a standard GPU cluster. Moreover, the features identified by SFA can serve as initial *collective variables* for enhanced sampling, providing flexibility in the choice of sampling strategy. Although slow feature analysis is used here, the protocol is general and can be adapted to alternative dimensionality reduction approaches, such as time-lagged autoencoders(40, 41) or principal component analysis(12, 42), depending on the motions of interest.

We further demonstrate how all-atom reconstruction of BioEmu generated conformational ensembles can be integrated with experimental cryo-EM density maps using a Bayesian reweighting framework to resolve the conformational ensemble and relative populations of multiple metastable states in apo GlyT1. GlyT1 is a membrane protein and key drug target for schizophrenia that exists in a dynamic equilibrium among three functionally relevant metastable states: occluded, inward, and outward. However, a direct comparison between BioEmu-augmented and rMSA-AF2-augmented molecular simulations reveal more limited sampling in the former (Figure 6 B, D), primarily due to reduced conformational coverage in the initial BioEmu-generated ensemble relative to rMSA-AF2. Notably, BioEmu-augmented simulations fail to capture flipping of Y62 in GlyT1, a functionally critical residue that acts as a “lid” and whose *flipping* is required for binding of the sarcosine-based inhibitor ALX-5407 and for driving the transition from the occluded to the inward state. Consistent with this role, mutation of Y62 to alanine significantly reduces glycine uptake activity, underscoring the functional importance of this side-chain motion(37). A similar deficiency in side-chain sampling is observed in apo PlmII, where flipping of the Trp41 side chain is a key motion governing the opening of a cryptic pocket. BioEmu does not explicitly model side-chain orientations; instead, side chains are grafted *post hoc* using external tools to generate all-atom representations of the predicted conformational ensemble. Because side-chain dynamics play a central role in regulating conformational transitions, as illustrated by both GlyT1 and PlmII, limited side-chain conformational diversity in these reconstructed ensembles may constrain the ability of subsequent physics-based molecular simulations to accelerate sampling of functionally relevant alternative states. Taken together, these observations challenge the growing assumption within the “generative AI for protein dynamics” community that prediction of backbone conformational heterogeneity alone is sufficient to capture protein dynamics.

**Figure 6.**
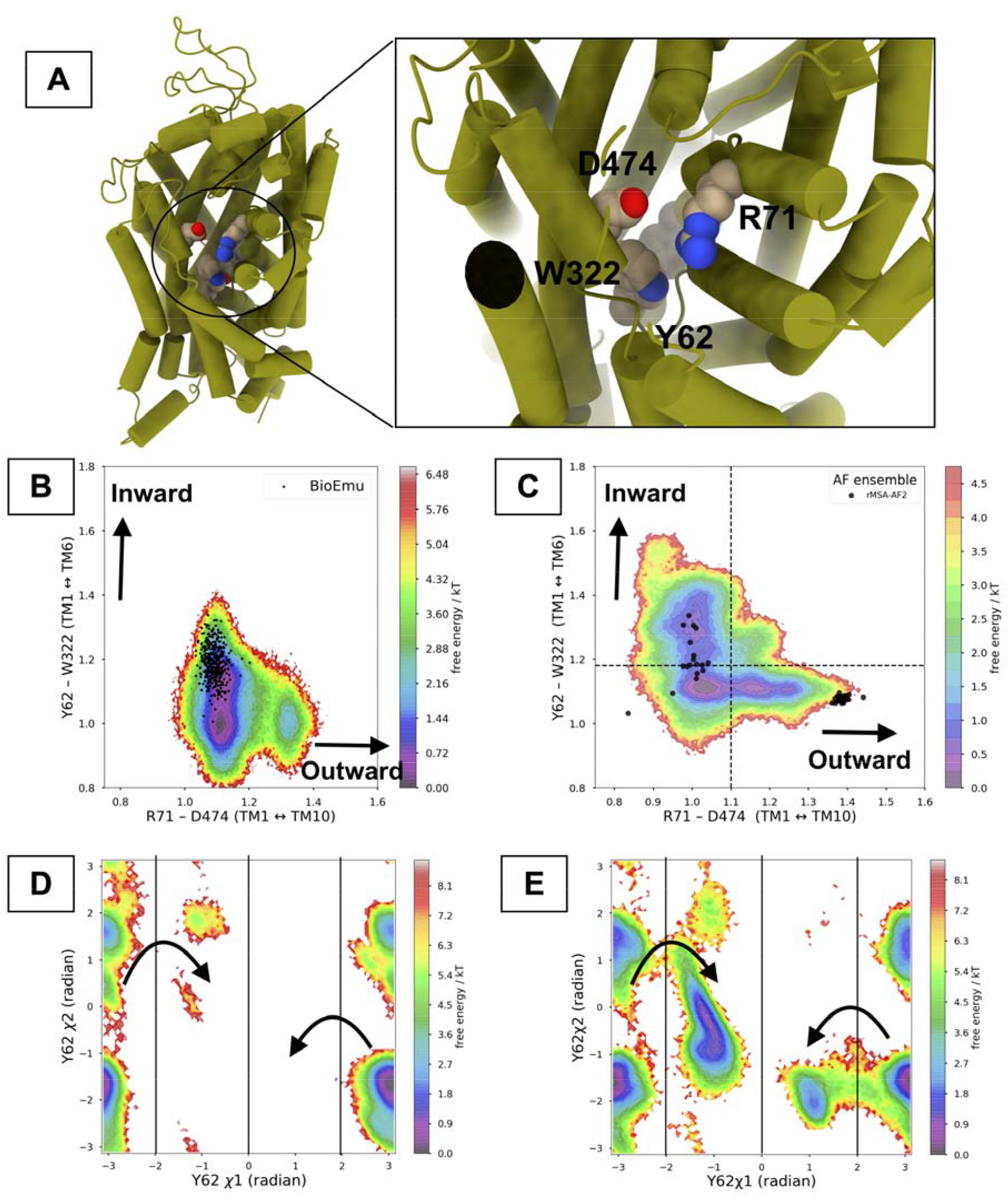
A pictorial representation of apo GlyT1 highlights conformational hotspots and key residues governing transitions between the inward, occluded, and outward states (A). Projections of Cα–Cα distances between residues defining TM1–TM6 and TM1–TM10 motions demonstrate more complete sampling of all three metastable states by rMSA-AF2 seeded molecular simulations compared to BioEmu seeded molecular simulations (B, C). MSM-weighted projection of the χ1 and χ2 dihedral angles of Y62 (a key determinant of GlyT1 function and conformational dynamics.) further reveal limited sampling in BioEmu augmented simulations (D) relative to rMSA-AF2 augmented simulations (E).

**Figure 7.**
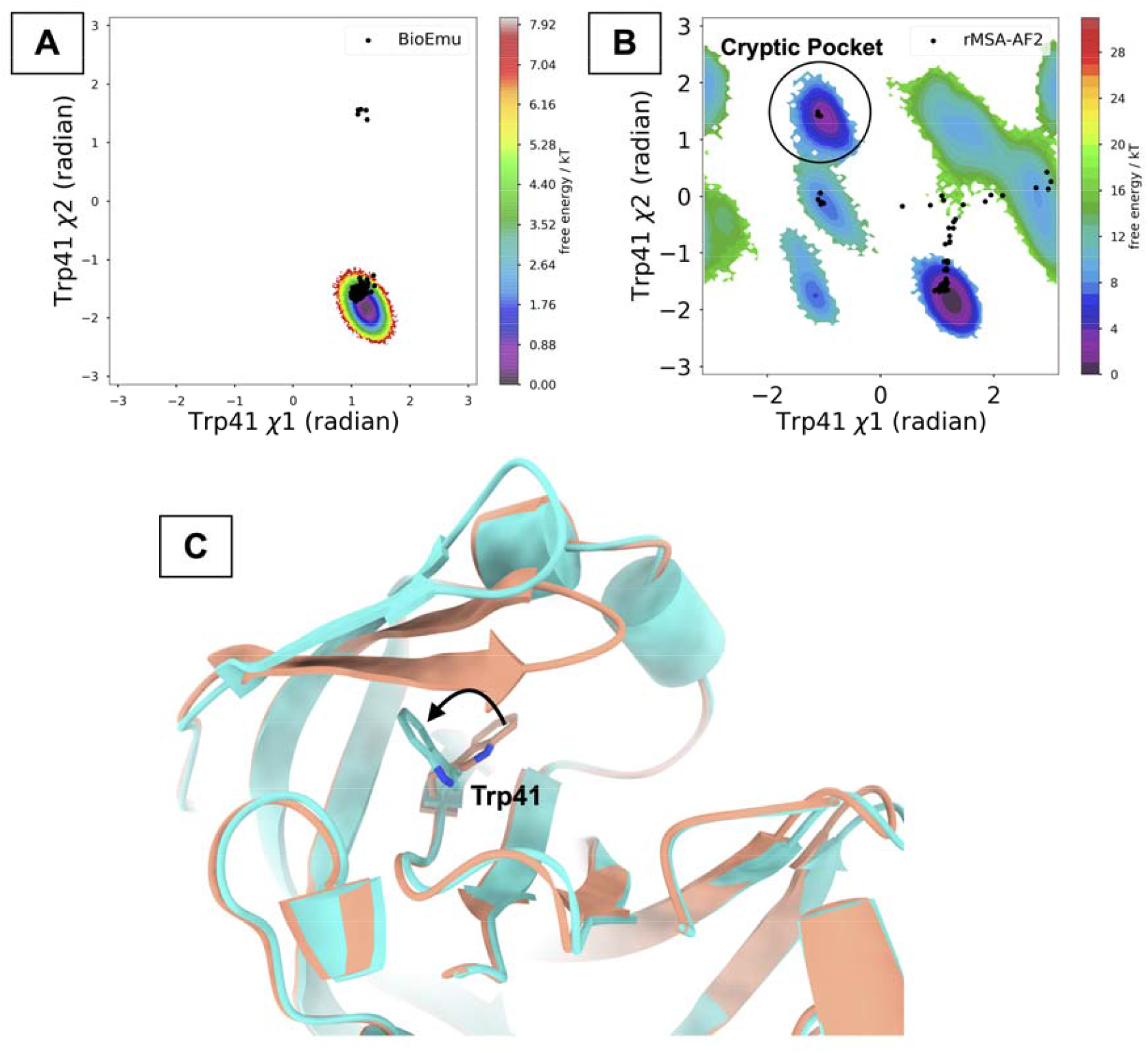
Boltzmann weighted free energy surface of side-chain conformational heterogeneity highlight the ability of rMSA-AF2 augmented molecular simulations (B) to capture cryptic pocket opening compared to BioEmu augmented molecular simulations (A). The initial conformational ensembles generated by BioEmu and rMSA-AF2 are shown as *black dots* and projected onto the corresponding free-energy surfaces, highlighting a wider conformational heterogeneity in rMSA-AF2. Superposition of the “open” (PDB: 2BJU) and “closed” (PDB: 1LF4) states highlights the flipping of the Trp41 side chain required for cryptic pocket opening (C).

A current limitation of this approach is that BioEmu v1.0 does not explicitly model protein– protein or protein–ligand interactions, limiting our ability to directly evaluate how these interactions reshape the conformational landscape. However, the overall framework is agnostic to interaction modality and can be readily extended to protein–protein and protein–ligand systems. As a practical recommendation, combining BioEmu generated ensemble with rMSA-AF2 as complementary priors can increase the likelihood of capturing broader conformational heterogeneity, particularly for systems where multiple metastable states coexist. Future efforts focused on fine-tuning BioEmu on class-specific molecular dynamics datasets will enable improved coverage of conformational ensembles in challenging protein classes that currently suffer from limited training data. More broadly, integrating generative AI models for protein ensemble generation with physics-based molecular simulations and single-molecule experiments (43)(44) provides a natural and scalable path forward to accurately capture protein dynamics and enable dynamics aware therapeutics discovery.

## Methods

### Ensemble generation

BioEmu(26) v1.0 (GitHub: https://github.com/microsoft/bioemu) was used to generate conformational ensembles comprising ∼500 structures for apo wild type CDK2, apo BRAF, apo V600E BRAF, apo human GlyT1 and apo PlmII. BioEmu automatically filters out structures with steric clashes or bond length violations from the generated ensemble; therefore, 500 represents the upper bound on the number of structures produced by this method. All-atom models were constructed by grafting side chains using H-packer (32) (GitHub: https://github.com/gvisani/hpacker) software.

### Feature selection and slow feature analysis

All Cα–Cα distances from the all-atom BioEmu generated conformational ensemble were converted into smooth, differentiable features using a continuous contact switching function (see *Bhakat et al*.(36)),

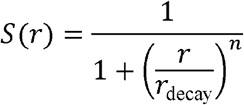

where *r* is the *Cα*−*Cα* distance, *r*_decay_ controls the transition length scale, and *n* sets the sharpness of the transition. Short distances (*r* « *r*_decay_) yield values close to 1 (formed contact), while long distances (*r* » *r*_decay_) yield values close to 0 (broken contact), providing a noise-robust, continuous measure of contact strength. We used *r*_decay_ = 0.90 nm and *n*= 6.

The Cα–Cα contacts were transformed using this function, yielding datapoints of *S*(*r*) ∈ [0,1]. To retain only dynamically informative contacts, we computed the minimum and maximum *S*(*r*) values for each contact across all trajectories and retained contacts satisfying

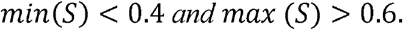

This filtering removes contacts that remain persistently formed or broken and selects contacts that undergo meaningful formation and disruption in the all-atom BioEmu ensemble. For interpretability, *S*(*r*) ≥ 0.6 corresponds to formed contacts, *S*(*r*) ≤ 0.4 to broken contacts, and intermediate values reflect transient contact fluctuations. The filtered contact set was used for subsequent dimensionality reduction.

The switching function transformed filtered distances were then used as input to Slow Feature Analysis (SFA), a dimensionality reduction technique for biomolecular simulations introduced by *Vats et al*.*(39)* in context of biomolecular data. SFA transforms the *J*-dimensional input signal **c**(*t*) into output signals *y*_*k*_ (*t*)= *g*_*k*_ (**c**(*t*)) by minimizing the temporal variation,

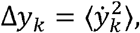

thereby extracting features that vary slowly over time. Applied to the filtered contact features, SFA identifies first two linear combinations of slowly fluctuating distance pairs that capture conformational hotspots from the all-atom BioEmu generated ensemble. K-means clustering (N=50) was subsequently performed on the first two slow features and subsequent cluster centers were saved in a PDB format for molecular simulation. SFA and clustering were performed using the MDML software package (GitHub: https://github.com/svats73/mdml/tree/main)

### Integration of BioEmu generated conformational ensemble with CryoPhold

For apo GlyT1, the cluster centers generated by SFA (N=50) were acted as a prior to enable Bayesian reweighting against inward (EMDB: EMD-37493; 3.35 Å), outward (EMDB: EMD-37494; 3.22 Å) and occluded (EMDB: EMD-37492; 2.58 Å) cryo-EM maps as describe in CryoPhold(25) (Github: https://github.com/strauchlab/cryoPhold).

### Molecular simulation

Representative structures from the SFA derived posterior ensemble were selected as starting points for unbiased molecular dynamics simulations. Each structure was prepared using the *tleap* module in Amber2022(43, 44) following the protocols described by *Meller et al*.*(24)* Proteins were parameterized with the AMBER ff14SB(45) force field, neutralized with counterions, and solvated in a truncated-octahedron TIP3P(46) water box with a minimum buffer of 10 Å from any protein atom to the box boundary. Energy minimization was performed in two stages: solvent and ions were first minimized with harmonic restraints on protein atoms (100 kcal mol□^1^ Å□^2^), followed by unrestrained minimization of the full system.

Amber topologies were converted to GROMACS format using ACPYPE(47), and all subsequent simulations were carried out in GROMACS 2022(48). Systems were heated from 0 to 300 K over 500 ps in the NVT ensemble with backbone heavy-atom restraints (500 kJ mol□^1^ nm□^2^), followed by 200 ps of unrestrained equilibration in the NPT ensemble at 300 K and 1 bar. Temperature was controlled using a velocity-rescale thermostat and pressure using a *Parrinello– Rahman*(49) barostat. Production simulations were performed in the NPT ensemble using a *2 fs* time step, a 1.0 nm cutoff for nonbonded interactions, particle-mesh Ewald(50) electrostatics, and LINCS(51) constraints on bonds involving hydrogen atoms.

Unbiased production simulations of 100 ns each were initiated from the SFA selected cluster centers, with trajectories saved every 10 ps, resulting in a cumulative simulation time of 5 μs for each protein system. rMSA-AF2 augmented molecular simulation data for GlyT1 and PlmII (total 8 μs for each protein system) were repurposed from Bhakat *et al(36) and Meller et al(24) respectively*.

Markov state models were constructed from the resulting trajectories using the PyEMMA(52) package. For kinases, the MSM was constructed using the distances highlighted in Figure 2 that distinguish DFG states. For apo GlyT1, the MSM was built using distances that discriminate among the inward, outward, and occluded states.

## Supporting information

Supporting Information revised

