## Supporting Information revised for "Accelerated sampling of protein dynamics using BioEmu augmented molecular simulation"

**^2^Division of Infectious Diseases, School of Medicine, Washington University in St. Louis, St. Louis, MO, United States**

 (SB)

 (EMS)

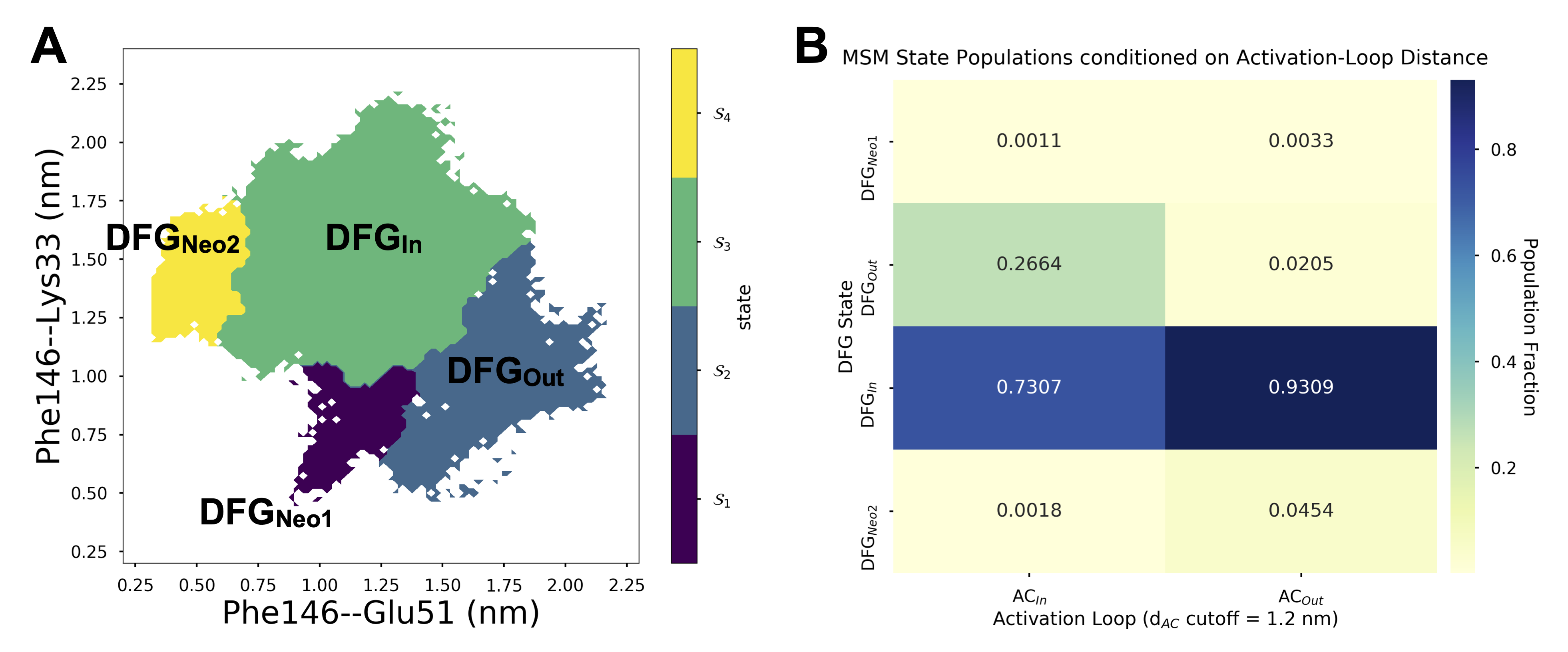

Figure S1. Overview of conformational macrostates captured by BioEmu-augmented molecular simulation of apo CDK2. **A:** Definition of macrostates using the PCCA+ algorithm, which separates canonical DFG-in and DFG-out states and identifies additional macrostates (denoted Neo1 and Neo2).
**B:** Markov State Model derived population fractions of extended (AC_out_) and folded (AC_in_) activation loop conformations within DFG states, illustrating the transition to the DFG-out state is accompanied by a shift toward the folded activation loop conformation.
